# Exploiting Heterogeneous Time Scale of Dynamics to Enhance 2D HETCOR Solid-State NMR Sensitivity

**DOI:** 10.1101/691220

**Authors:** Rongchun Zhang, Yusuke Nishiyama, Ayyalusamy Ramamoorthy

## Abstract

Multidimensional solid-state NMR spectroscopy plays a significant role in offering atomic-level insights into molecular systems. In particular, heteronuclear chemical shift correlation (HETCOR) experiments could provide local chemical and structural information in terms of spatial heteronuclear proximity and through-bond connectivity. In solid state, the transfer of magnetization between heteronuclei, a key step in HETCOR experiments, is usually achieved using cross-polarization (CP) or INEPT (insensitive nuclei enhanced by polarization transfer) depending on the sample characteristics and magic-angle-spinning (MAS) frequency. But, for a multiphase system constituting molecular components that differ in their time scales of mobilities, CP efficiency is pretty low for mobile components because of the averaging of heteronuclear dipolar couplings whereas INEPT is inefficient due to the short T_2_ of immobile components and can be non-selective due to strong proton spin diffusion for immobile components especially under moderate spinning speeds. Herein, in this study we present two 2D pulse sequences that enable the sequential acquisition of ^13^C/^1^H HETCOR NMR spectra for the rigid and mobile components by taking full advantage of the abundant proton magnetization in a single experiment with barely increasing the overall experimental time. In particular, the ^13^C-detected HETCOR experiment could be applied under slow MAS conditions, where a multiple-pulse sequence is typically employed to enhance ^1^H spectral resolution in the indirect dimension. In contrast, the ^1^H-detected HETCOR experiment should be applied under ultrafast MAS, where CP and transient heteronuclear nuclear Overhauser effect (NOE) polarization transfer are combined to enhance ^13^C signal intensities for mobile components. These pulse sequences are experimentally demonstrated on two model systems to obtain 2D ^13^C/^1^H chemical shift correlation spectra of rigid and mobile components independently and separately. These pulse sequences can be used for dynamics difference based spectral editing and resonance assignments. Therefore, we believe the proposed 2D HETCOR NMR pulse sequences will be beneficial for the structural studies of heterogeneous systems containing molecular components that differ in their time scale of motions for understanding the interplay of structures and properties.

## 1. Introduction

Solid-state NMR spectroscopy is a powerful analytical tool that is widely used for studying molecular structures and dynamics of a variety of solids including membrane proteins,[1–5] multiphase polymers[6–8], bone materials,[9–11] amyloids[12–17], and nanocomposites[18–21]. The benefits of multidimensional NMR techniques to obtain higher resolution and higher degree of structural and dynamic information have been well utilized in numerous applications, among which 2D homonuclear (HOMCOR)[22–25] and heteronuclear (HETCOR)[26–31] chemical shift correlation experiments are most widely employed and have become indispensable for structural studies of proteins.[32–39]

With the rapid development of ultrafast magic-angle-spinning (MAS) probe technology, ^1^H spectral resolution can be dramatically enhanced due to efficient suppression of ^1^H-^1^H dipolar couplings,[40, 41] which allow for ^1^H/^1^H chemical shift correlation [42, 43] to probe the proximities of proton atoms in solids. Most importantly, proton-detection under ultrafast MAS conditions is employed to remarkably improve the signal-to-noise (S/N) ratio of multidimensional experiments.[44–52] In these experiments involving low-*γ* nuclei, cross polarization (CP) [53, 54] is generally employed to enhance the S/N of low-*γ* nuclei, where the polarization transfer efficiency largely relies on the dipolar couplings between proton and low-*γ* nuclei. As a result, polarization transfer can be difficult if heteronuclear dipolar couplings are greatly averaged out by molecular motions, as often encountered for the mobile components in semi-solids or multiphase heterogeneous systems. To enhance the S/N of mobile components, we recently demonstrated a method for acquiring 1D ^13^C MAS spectrum by combining CP and NOE (transient heteronuclear nuclear Overhauser effect) or RINEPT (refocused insensitive nuclei enhanced by polarization transfer), dubbed as CP-NOE and CP-RINEPT, respectively.[55] For a multi-component system, the ^13^C signals from rigid components are enhanced by CP, while the signals from mobile components are enhanced by either heteronuclear NOE or RINEPT. In addition, the CP-RINEPT experiment enables the acquisition of two separate spectra, each of which contains signals from either rigid or mobile components alone.

In this study, we propose two 2D HETCOR pulse sequences utilizing CP-NOE or CP-RINEPT sequences to enhance the ^13^C signals of mobile components. The ^13^C-detected HETCOR experiment can be used under slow MAS speeds, where a multiple-pulse sequence is employed to enhance the ^1^H resolution in the indirect dimension. On the other hand, the ^1^H-detected HETCOR experiment should be performed under very fast MAS, where the sensitivity is enhanced by the combination of CP and heteronuclear NOE via ^1^H detection. Compared to the regular 2D HETCOR experiment, the proposed method offers the acquisition of two separate 2D HETCOR spectra for rigid alone and mobile alone components of a multiphase system with barely increasing the overall experimental time. Herein, the 2D HETCOR spectra obtained for rigid and mobile components are denoted as Rigid-HETCOR and Mobile-HETCOR, respectively. The benefits and possible limitations of the two proposed methods are discussed within the context of their applications to study polymer blend systems.

## 2. Radio-frequency pulse schemes

The radio-frequency (RF) pulse sequences demonstrated in this study are shown in Figure 2. The ^13^C-detected 2D HETCOR NMR experiment (Figure 2a) is designed for a moderate MAS frequency. In this sequence, during the *t*_1_ period, the transverse ^1^H magnetization is allowed to evolve under Frequency-switched Lee-Goldburg (FSLG)[56–59] pulse sequence that is used to suppress ^1^H-^1^H dipolar couplings. While the transverse ^1^H magnetization from rigid components is transferred to ^13^C using CP, those from the mobile components are kept spin-locked during CP. During the first signal acquisition period for the Rigid-HETCOR spectrum, high power continuous-wave (CW) RF decoupling is applied to achieve high spectral resolution and also to retain the transverse ^1^H magnetization by spin-locking for the subsequent RINEPT-based magnetization transfer. While the RINEPT sequence transfers the transverse ^1^H magnetization from mobile components, it suppresses the ^1^H magnetization from rigid components due to their short T_2_ relaxation, and thus enables acquiring Mobile-HETCOR spectrum through the second t_2_ acquisition period.

**Figure 1.**
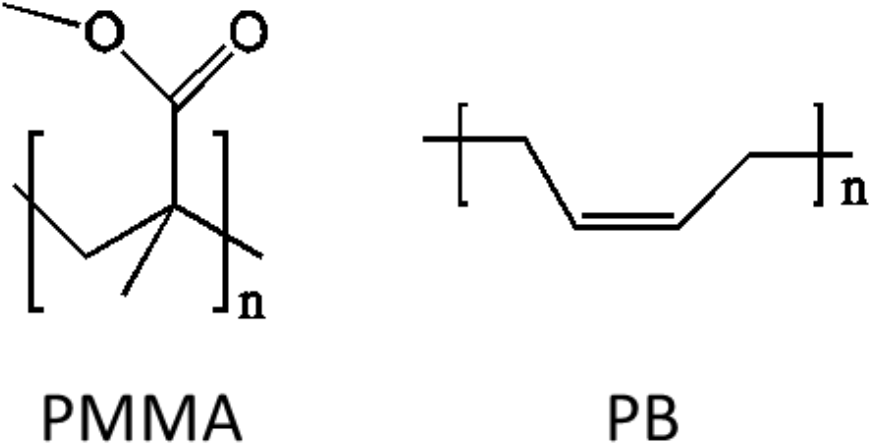
Chemical structures of PMMA and PB used in this study.

**Figure 2.**
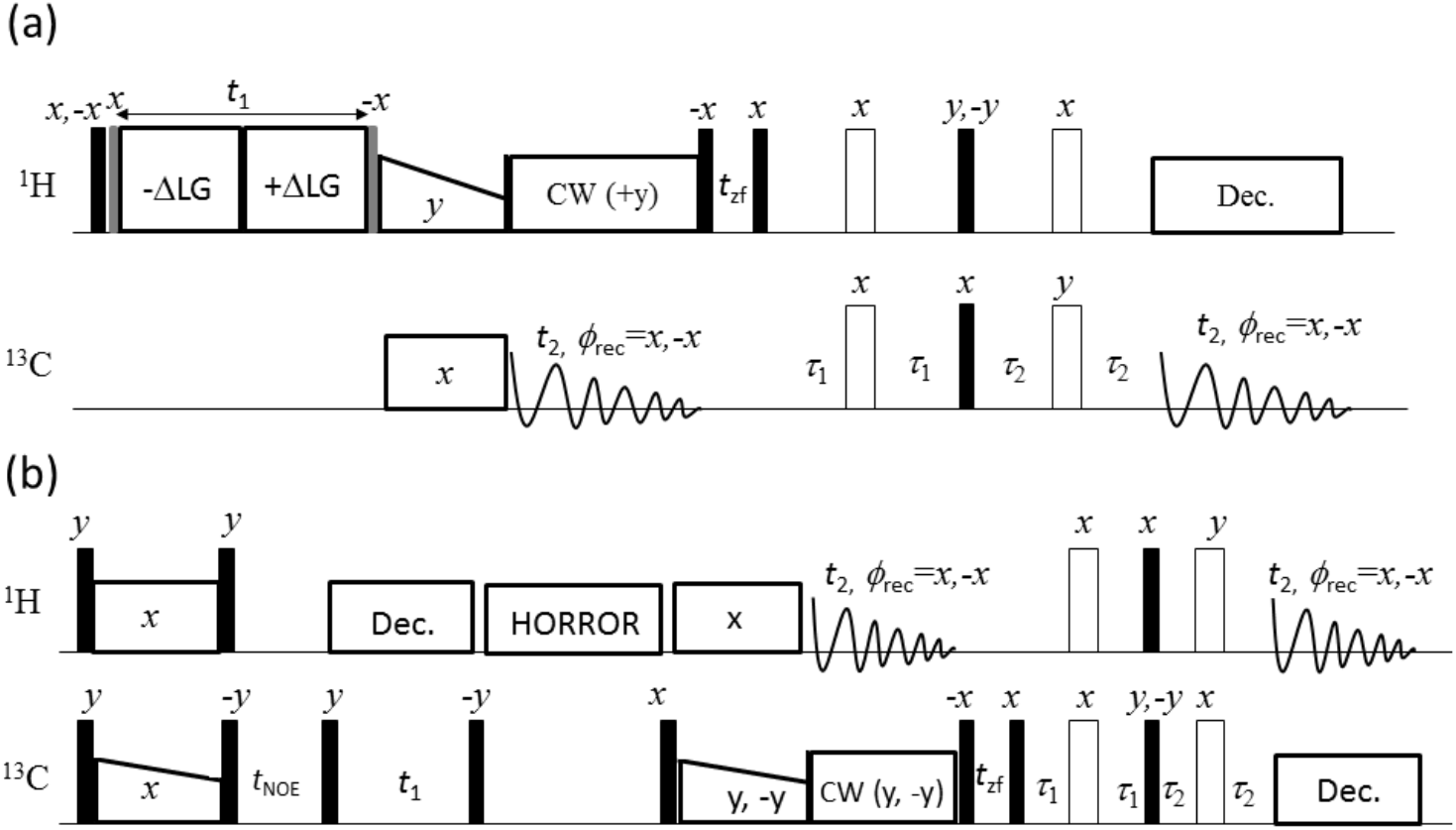
2D ^13^C-detected (a) and ^1^H-detected (b) 2D HETCOR NMR pulse sequences applied under slow and very fast MAS, respectively. The black solid and blank rectangles indicate 90° and 180° pulses, respectively. The grey solid rectangles are 35.3° RF pulses applied before and after the incrementable *t*_1_ period in (a). The 35.3° pulse before the *t*_1_ period flips the transverse magnetization to a plane transverse to the effective field of the FSLG sequence during the *t*_1_ period, whereas the 35.3° pulse after the *t*_1_ period flips the magnetization to plane transverse to the B_o_ field.

The ^1^H-detected HETCOR NMR experiment under fast-MAS is shown in Figure 2b. After the initial CP period, the transverse ^13^C magnetization is flipped to the +z direction for storage, while the ^1^H magnetization is flipped to -z direction to enable the transient ^1^H→^13^C heteronuclear NOE during *t*_*NOE*_. Because the NOE based polarization transfer relies on molecular motions, only the signals from the mobile components are enhanced in intensity, while the signals from the rigid components are retained due to their long ^13^C spin-lattice relaxation (*T*_*1*_) time. In this study, the NOE mixing time (*t*_*NOE*_) was generally set around 1s to maximize the NOE-based polarization transfer while at the same time minimizing the spin-lattice relaxation effect on the loss of signals from the rigid component. After NOE mixing period, a 90° pulse is applied in the ^13^C RF channel to allow ^13^C evolution during t_1_ and then the magnetization is stored along the z axis by the following 90° pulse for both mobile and rigid components. The HORROR sequence [60] is applied to remove any residual ^1^H magnetization before ^13^C→^1^H CP. The ^13^C magnetization from the rigid components is transferred to _1_H by the second CP for the first signal acquisition during *t*_2_ (to obtain the 2D Rigid-HETCOR spectrum). On the other hand, the ^13^C magnetization from the mobile components is spin-locked during the ^13^C→^1^H CP and subsequent CW decoupling periods, and then transferred to protons by RINEPT for the second signal acquisition during *t*_2_ (Mobile-HETCOR). During signal acquisitions of rigid and mobile components ^13^C nuclei can be decoupled as indicted in the pulse sequence.

## 3. Experimental Details

### 3.1. Materials

The hydroxyl-terminated 1,4-polybutadiene (PB) oligomers were purchased from Qilu Ethylene Chemical and Engineering Co. Ltd. (China). All the other samples were purchased from Aldrich Co., and used as received without any further purification. Molecular weights of poly(methyl methacrylate) (PMMA) and PB are 550,000 g/mol and 4,200 g/mol, respectively. The glycine/adamantane mixture was prepared with a 1:1 weight ratio. The PMMA/PB blend was prepared by mechanically mixing PMMA and PB in a 6:1 weight ratio.

### 3.2. Solid-State NMR Experiments

The ^13^C-detected HETCOR NMR experiment (Figure 2a) was performed on a Varian 400 MHz VNMRS solid-state NMR spectrometer equipped with a 5 mm triple-resonance MAS probe. The spinning speed was actively controlled at 8 kHz with a MAS speed controller. The ^1^H and ^13^C resonance frequencies were 400.11 MHz and 100.62 MHz, respectively. The 90° pulse widths were 3.2 μs for ^1^H and 4.4 μs for ^13^C, respectively. The effective FSLG decoupling field strength was 95.7 kHz, corresponding to a frequency offset of 55.3 kHz. Ramped-CP with a 5% ramp on the ^1^H channel was used to transfer ^1^H magnetization to ^13^C. The RINEPT evolution times, *τ*_1_ and *τ*_2_, were set to 1.4 ms and 1.0 ms, respectively. The CP contact time was set as 100 μs in order to obtain only the bonded ^13^C-^1^H chemical shift correlation in the final 2D HETCOR spectra.

The ^1^H-detected HETCOR (Figure 2b) NMR experiments of PMMA/PB blend were performed on an Agilent VNMRS 600 MHz solid-state NMR spectrometer equipped with a triple-resonance 1.2 mm MAS probe. The MAS speed was actively controlled at 60 kHz. The 90° pulse width was set to 2.0 μs for both ^1^H and ^13^C nuclei. Ramped-CP with a 17% ramp on the ^13^C channel was used for transferring the magnetization from ^1^H to ^13^C and from ^13^C to ^1^H in Figure 2b. The RINEPT evolution times, *τ*_1_ and *τ*_2_, were both set to 1.0 ms. The contact times for the first and second CP were 2 and 0.4 ms, respectively. The NOE mixing time (*t*_NOE_) and recycle delay were 1 and 3 s, respectively. The ^1^H-detected HETCOR experiment on glycine/adamantane mixture was performed under 100 kHz MAS on a 900 MHz ECZ900R solid-state NMR spectrometer equipped with a 0.75 mm double resonance ultrafast MAS probe (JEOL Resonance Inc). The 90° pulse widths were 0.6 and 1.2 μs for ^1^H and ^13^C nuclei, respectively. The contact times for the first and second CP were 0.5 and 0.2 ms, respectively. Both the RINEPT evolution times, *τ*_1_ and *τ*_2_, were set to 1.0 ms and a 2.5 s recycle delay was used. The NOE mixing time (*t*_NOE_) was 2 s.

For both ^13^C and ^1^H-detected HETCOR experiments shown in Figure 2(a and b), a z-filter of ~3 ms was inserted right after the first signal acquisition period to remove the residual magnetization in the transverse plane. The ^13^C chemical shifts in all spectra are referenced with respect to TMS using the low-frequency peak of adamantane (*δ*_iso_ = 29.4 ppm) as a secondary reference. Two 2D HETCOR spectra can be obtained from a single experiment, which are denoted as Rigid-HETCOR and Mobile-HETCOR spectra.

## 4. Results and Discussion

Molecular motion plays a critical role in most molecular systems, which may significantly affect the physical, chemical and functional properties of a material.[61–63] For example, the mobile components in a multiphase polymer system can significantly increase the toughness of the material, and thus lead to an enhancement of stretching extensibility. Although such polymeric materials are not amenable for atomic-resolution studies for most analytical techniques, they can be investigated at atomic-resolution using a variety of solid-state NMR techniques that have no limitation on the molecular size and nature of the material. However, in a typical CP based MAS experiment, the signals of mobile components are greatly compromised or even lost due to the their weak ^1^H-^13^C heteronuclear dipolar couplings, rendering the traditional CP-based HETCOR experiment incapable to detect ^1^H/^13^C chemical shift correlations of mobile components. In order to address this issue, herein we have successfully demonstrated two HETCOR pulse sequences (Figure 2) suitable for measurements under slow and very fast MAS conditions, enabling the acquisition of 2D HETCOR NMR spectra of rigid and mobile components in a single experiment without a significant increase in the overall experimental time by making use of a single recycle delay. To demonstrate the efficacy of the proposed pulse sequences, a glycine and adamantane mixture was first used as a model system before applying the techniques on a complex polymer blend. Glycine is a rigid crystalline small molecule with α or γ form stable crystallites, while adamantane can be considered as a mobile component since its fast molecular rotations significantly suppress intramolecular ^1^H-^1^H dipolar interactions. It should be noted that residual ^1^H-^1^H and ^1^H-^13^C intermolecular interactions remain in adamantane since it is a plastic crystal. Indeed, the Rigid-HETCOR spectrum shown in Figure 3a obtained using the ^13^C-detected HETCOR pulse sequence (Figure 2a) gives both glycine and adamantine signals even with a very short contact time of 0.1 ms. This is because the presence of ^1^H-^1^H homonuclear dipolar interaction speeds up the CP ^1^H-^13^C build-up in addition to residual intermolecular ^13^C-^1^H dipolar couplings in adamantane. However, in the Mobile-HETCOR spectra shown in Figure 3b, only adamantane signals were observed due to the use of RINEPT sequence, where the rigid glycine signals are suppressed by the fast spin-spin relaxation during the evolution periods (τ_1_ and τ_2_). These spectra also demonstrate the use of FSLG in the *t*_1_ (i.e. ^1^H) dimension to obtain narrow proton peaks under a slow spinning speed.

**Figure 3.**
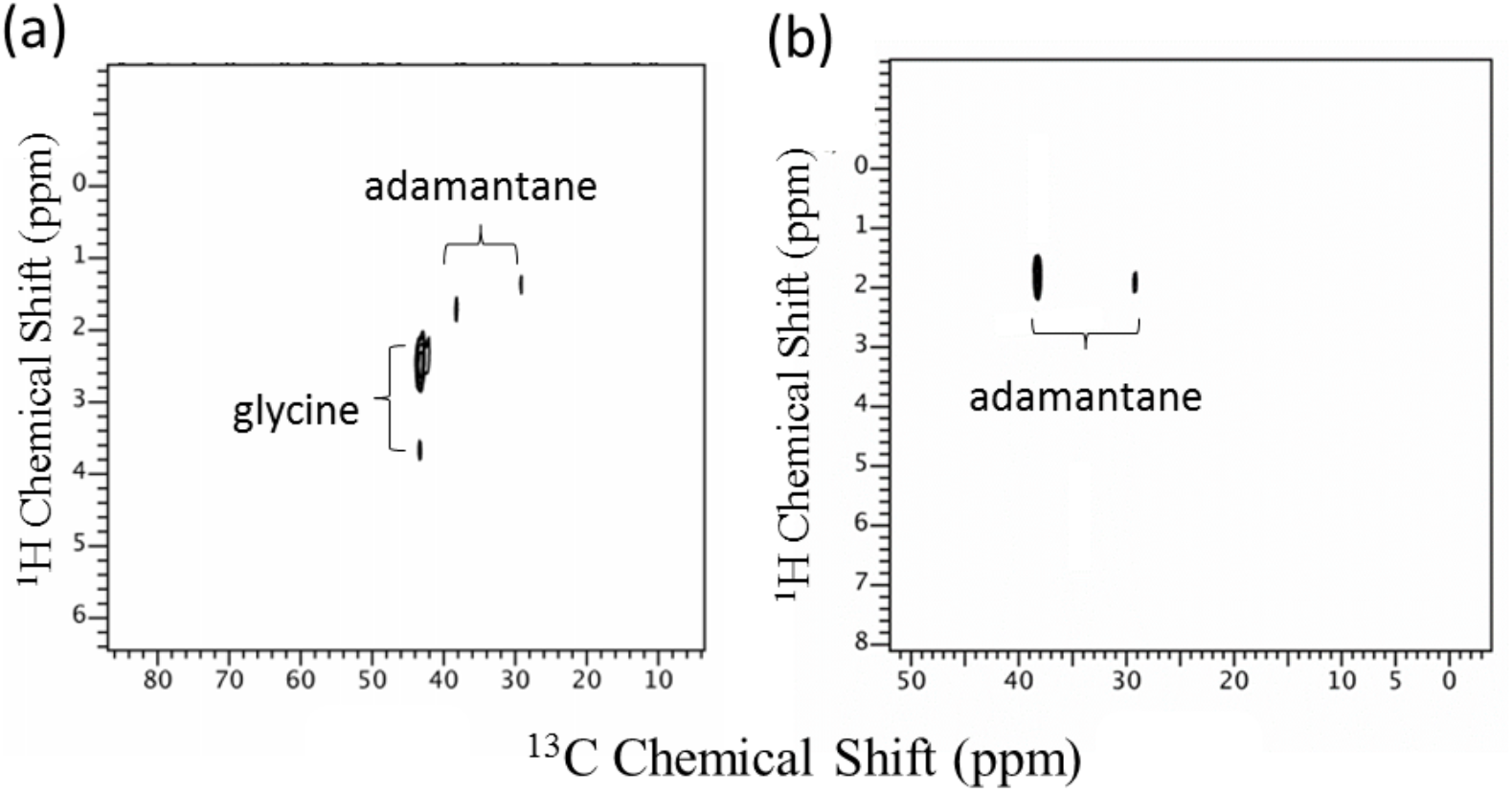
^13^C-detected HETCOR spectra of a glycine/adamantine mixture under 8 kHz MAS: Rigid-HETCOR (a) and Mobile-HETCOR (b) spectra were obtained from a single experiment using the pulse sequence shown in Figure 2a. A CP contact time of 0.1 ms, and the RINEPT evolution times, *τ*_1_ and *τ*_2_, of 1.4 ms and 1.0 ms, respectively, were used.

Using the pulse sequence shown in Figure 2b, ^1^H-detected 2D HETCOR spectrum obtained from glycine/adamantine mixture under 100 kHz MAS is shown in Figure 4. For a direct comparison, the Rigid-HETCOR and Mobile-HETCOR spectra are superimposed. We did not observe any adamantane signal in the Rigid-HETCOR spectrum at 100 kHz MAS, although the second CP contact time of 0.2 ms is slightly longer than that used for experiments at 8 kHz MAS. This is because the CP build-up curve is dominated by the weak intermolecular heteronuclear dipolar interactions due to the suppression of ^1^H-^1^H dipolar interactions under 100 kHz MAS. Because of this, signals from glycine and adamantane are well separated, and two independent HETCOR spectra corresponding to that of glycine and adamantane are obtained in a single experiment. It should be noted that we did not observe two CH_2_ proton peaks for glycine in the Rigid-HETCOR spectrum, since the glycine crystallite is in the γ form instead of the α form as used for the ^13^C-detected HETCOR experiment under a slow spinning speed. The capabilities of ultrafast MAS as well as the multiple-pulse sequence in resolving the two proton peaks of α form glycine crystallites have been well demonstrated in multiple studies.[64, 65] In this study, we have demonstrated the performance of ^13^C-detected and ^1^H-detected HETCOR pulse sequences to obtain 2D ^13^C/^1^H HETCOR spectra of rigid and mobile components separately in a single experiment.

**Figure 4.**
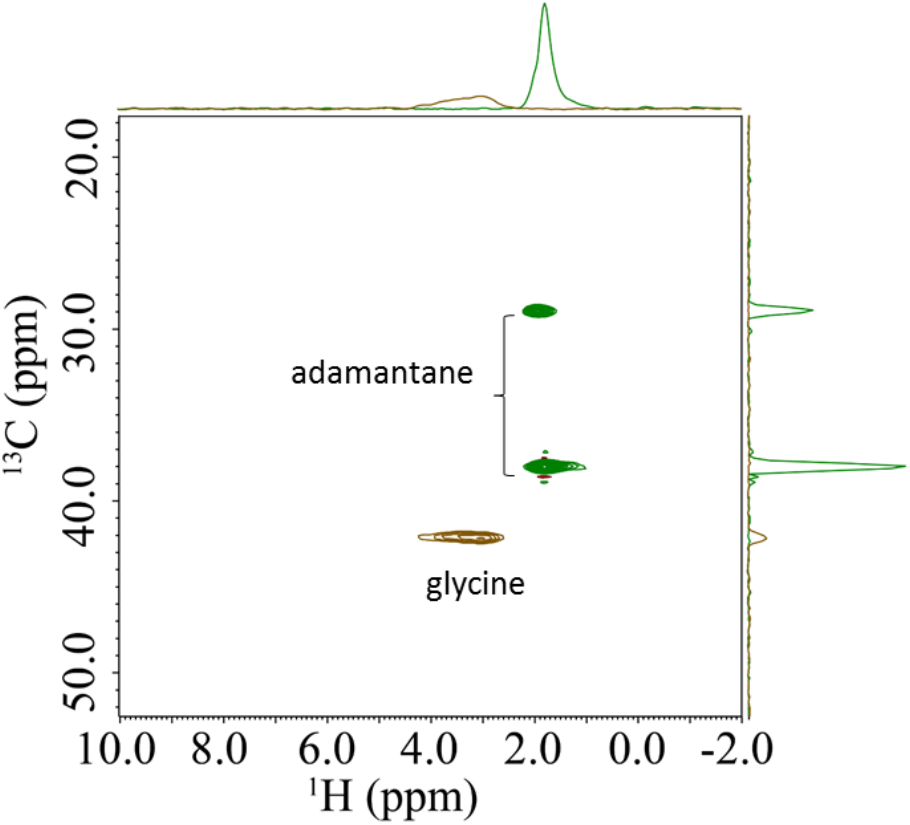
^1^H-detected HETCOR spectra of glycine/adamantine mixture under 100 kHz MAS. The superimposed Rigid-HETCOR (brown) and Mobile-HETCOR (green) spectra were obtained from a single experiment using the pulse sequence shown in Figure 2b. 0.5 ms (first CP) and 0.2 ms (second CP) contact times, and 2 s NOE mixing time (*t*_NOE_) were used.

Polymer blend has many technological applications and is widely used in industry since the merits of two different bulk materials can be combined and the end-use properties (*eg*. mechanical and rheological) can be optimized.[66–69] In this study, we further demonstrated the robust performance of the proposed pulse sequences on polymer blend PMMA/PB; their chemical structures are shown in Figure 1. PMMA is a semi-crystalline polymer with glass transition (*T*_g_) and melting (*T*_m_) temperatures both above 100 °C, whereas PB is a viscous liquid with a *T*_g_ of ~ −90 °C. Therefore, in the PMMA/PB blend, PMMA is considered as the rigid and PB is the mobile components. It was previously shown that in such a polymer blend system with a significant mobility contrast for the components, the ^13^C signals of PB cannot be detected by CP,[55] because the ^13^C-^1^H dipolar couplings have been completely averaged out by the fast molecular motion of PB at room temperature. However, the ^13^C signals of such mobile components can be detected by the RINEPT sequence or transient heteronuclear NOE.[55] Herein, using a combination of CP and RINEPT and/or NOE, ^13^C-detected and ^1^H-detected HETCOR (Figure 2) experiments can be used to obtain the ^13^C/^1^H HETCOR spectra of such polymer blend, where the HETCOR spectra of rigid and mobile components can be separately obtained in a single experiment. The 2D spectra obtained from the PMMA/PB blend are shown in Figures 5 and 6. The Rigid-HETCOR spectrum exhibits signals only from rigid PMMA, while Mobile-HETCOR spectrum displays the mobile PB signals. It may be noted that the proton peaks of PB signals in Mobile-HETCOR spectrum appears to be very broad, which can be attributed to the presence of different conformation (trans- and cis-1,4-units, >80%), the vinyl-1,2-units and end-groups in PB chains[70]; and also partly due to truncation of signal as the longest *t*_1_ of ~10 ms may be insufficient and the use of an exponential line broadening factor of 100 Hz in *t*_1_ during 2D Fourier transformation to reduce the noise. However, the trans- and cis-1,4-units in PB still dominate the signals, and thus clear ^13^C/^1^H chemical shift correlations can be observed for the dominating cis- and trans-1,4-PB chains.

**Figure 5.**
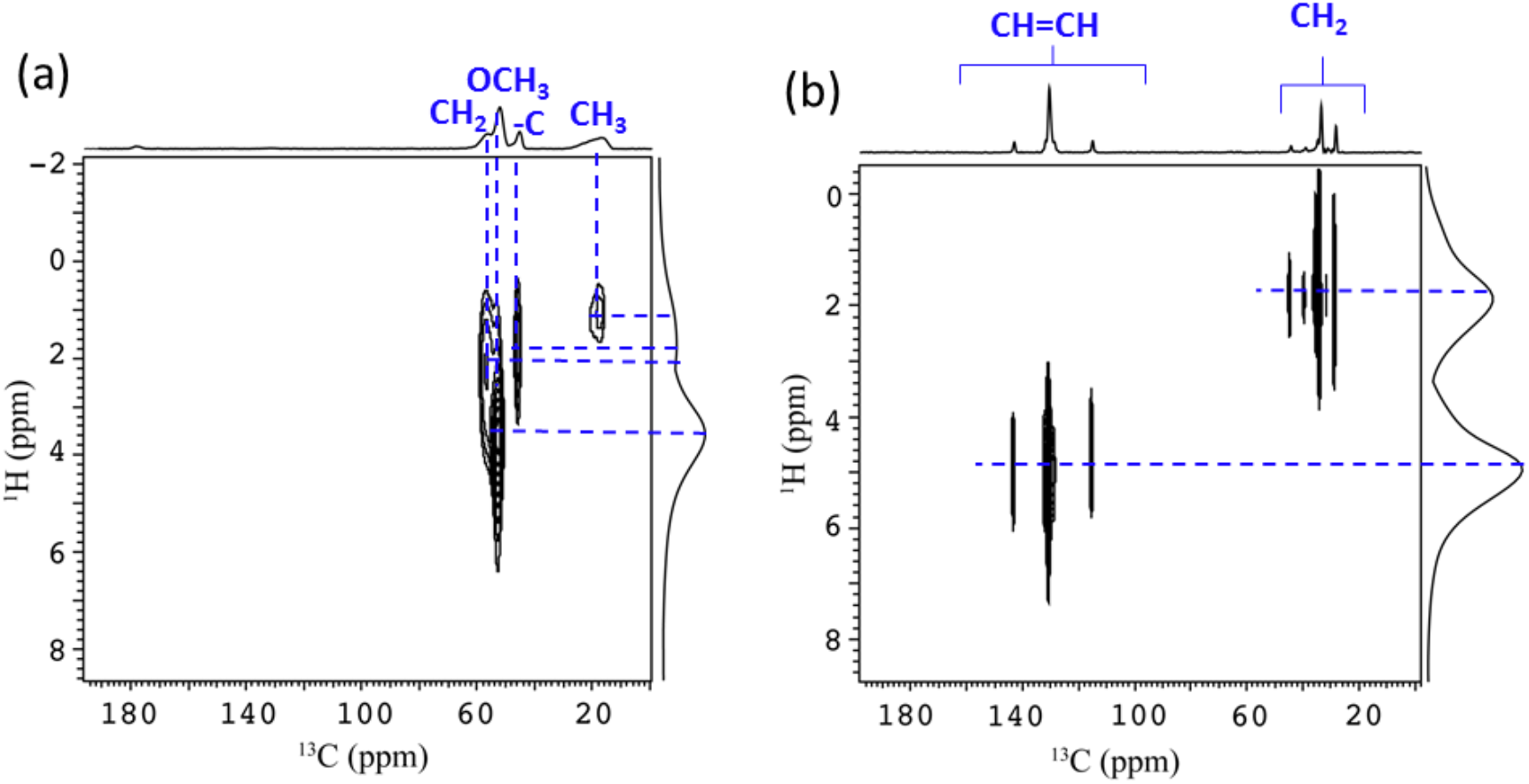
^13^C-detected HETCOR spectra of PMMA/PB blend obtained under 8 kHz MAS: Rigid-HETCOR (a) and Mobile-HETCOR (b) spectra were obtained from a single experiment using the pulse sequence shown in Figure 2a. 0.1 ms CP contact time and the RINEPT evolution times, *τ*_1_ and *τ*_2_, of 1.4 ms and 1.0 ms, respectively, were used.

**Figure 6.**
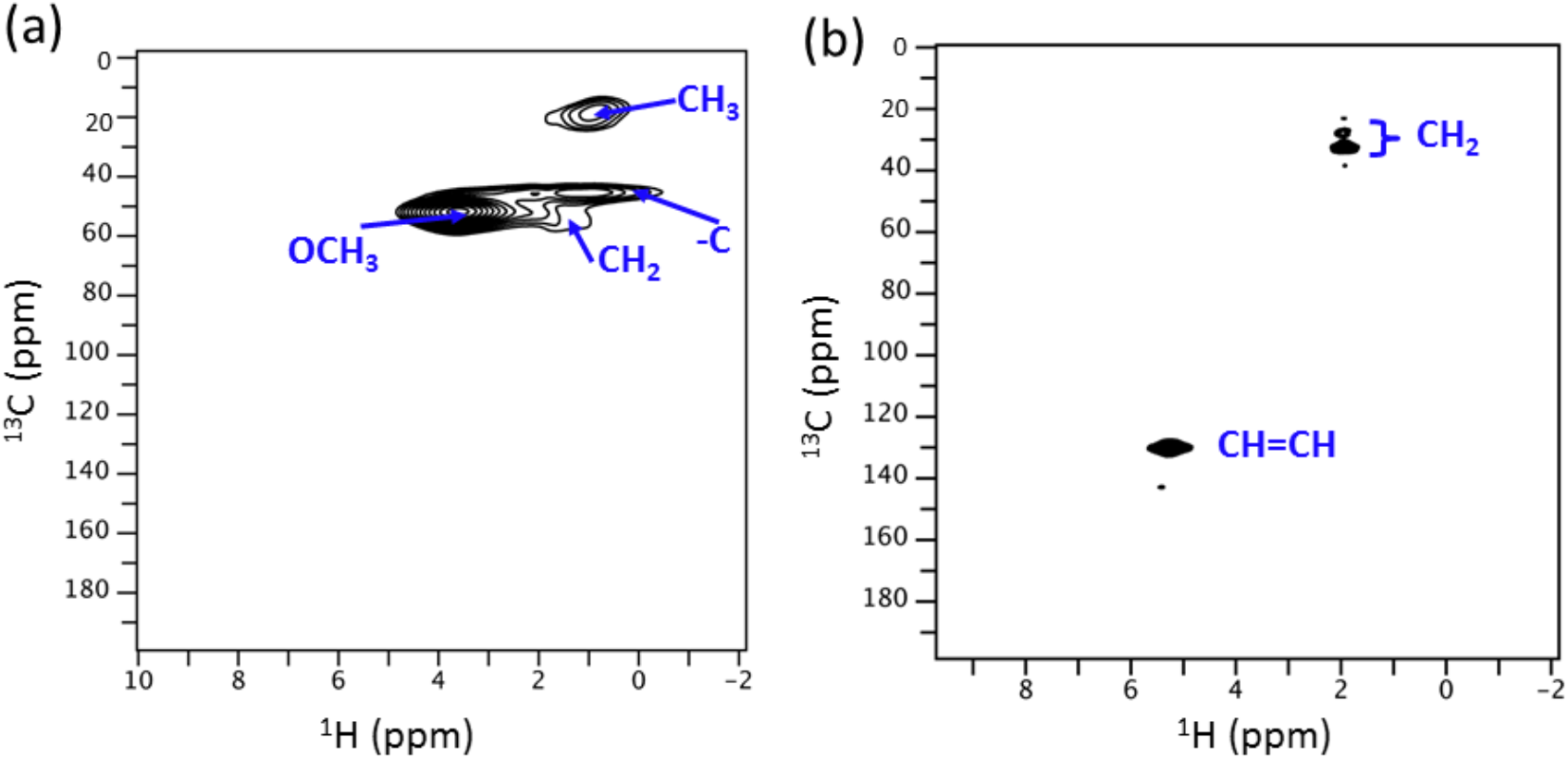
^1^H-detected CR-HETCOR spectra of PMMA/PB blend obtained under 60 kHz MAS: Rigid-HETCOR (a) and Mobile-HETCOR (b) spectra were obtained simultaneously from a single experiment using the pulse sequence shown in Figure 2b. First and second CP contact times of 2 and 0.4 ms, respectively, and 0.5 s NOE mixing time (*t*_NOE_) were used.

In the ^1^H-detected 2D Mobile-HETCOR spectra (Figure 6b) obtained using the ^1^H-detected HETCOR pulse sequence (Figure 2b), signals from the minor components in PB (vinyl-1,2-units, end-groups, *etc*.) are not observed due to the small sample volume in the rotor (~1.2 μL). In addition, the direct detection of proton signals after the RINEPT sequence also results in higher signal intensity as a much longer acquisition time (a few hundreds of milliseconds) can be used along with/without low-power decoupling on the ^13^C RF channel. For the two pulse sequences, the continuous-wave (CW) decoupling has to be applied during the signal acquisition after CP in order to keep the transverse magnetization of mobile components for the subsequent polarization transfer by RINEPT. For the ^13^C-detected HETCOR experiment (Figure 2a), CW decoupling might potentially compromise ^13^C spectral resolution of rigid components. However, it should be noted that in the multiphase solids, the mobility of rigid components is generally enhanced by the mobile components; thus CW decoupling with high RF strength can still provide a reasonable spectral resolution as demonstrated in our previous study.[55] Notably, the ^13^C-detected HETCOR experimental time is nearly the same as that of the traditional ^13^C-detected CP-based HETCOR experiment. However, for the ^1^H-detected HETCOR experiment the increase in experimental time, compared to the traditional ^1^H-detected CP-based HETCOR experiment, is mostly due to the NOE mixing time (~ 1s) in each scan.

Indeed, in terms of shortening experimental time as well as enhancing signal sensitivity of multi-dimensional solid-state NMR experiments, several approaches and strategies have been proposed to enable multiple acquisition in a single scan, either through elegant designs of pulse sequences or with the use of multiple receivers.[71–76] In those multi-acquisition experiments, the ^13^C or ^15^N magnetizations are generally re-utilized as the source for polarization transfer after the first acquisition period, because the ^13^C or ^15^N magnetization is easy to store due to the very long spin-lattice relaxation time (T_1_) of these nuclei. However, the use of residual proton magnetization is much more challenging due to its short T_1_ and T_1ρ_, but greatly beneficial for the signal enhancement of low-*γ* nuclei as demonstrated by our work here and previous studies.[55, 77]

## 5. Conclusions

In multi-component solids with mobility contrast, the ^13^C signals of mobile components are undetectable in the regular CP-based HETCOR experiment due to insufficiency of CP for polarization transfer, since the ^1^H-^13^C dipolar couplings are averaged out by the fast molecular motions. Herein, we have proposed two novel HETCOR pulse sequences (^13^C- and ^1^H-detected HETCOR) suitable for measurement under slow and ultrafast MAS conditions by fully utilizing the abundant proton magnetization for polarization transfer. From a single experiment, two HETCOR spectra, through the polarization transfer based on CP and RINEPT, can be obtained corresponding to that of rigid and mobile components, respectively. In the ^1^H-detected HETCOR experiment, transient heteronuclear NOE was well combined with CP to significantly enhance the ^13^C magnetization of mobile components in the indirect dimension, and thus enables the subsequent ^13^C→^1^H polarization transfer with RINEPT sequence. While two HETCOR spectra can be obtained from such a single experiment, the experimental time does not increase significantly, as compared to the single conventional CP and RINEPT-based HETCOR experiment. Thus, the total experimental time was significantly reduced when compared to the total time needed to measure two conventional CP and RINEPT-based HETCOR experiment to obtain HETCOR spectra for rigid and mobile components, respectively. In addition to the significant reduction of data collection time, the separation of signals from mobile and rigid components can also be potentially used for resonance assignment in a wide array of multiphase heterogeneous systems. Thus, we believe the proposed sequences here will greatly beneficial for the structural elucidation of complex systems, such as biomaterials, polymer blends, proteins, supramolecular assemblies and so on. For example, atomic-resolution investigation of the ubiquitous self-assembly process underlying the formation of nanostructures including amyloid fibrils, using the proposed 2D HETCOR pulse sequences, would enable selective monitoring of fast-tumbling monomers and small oligomers even in the presence of large aggregates or fibers.[78]Such experiments can shed light on the nucleation of small oligomeric amyloid intermediates that have been shown to be more toxic in the amyloid diseases like Alzheimer’s Disease and type II diabetes.

## 6. Acknowledgments

We acknowledge the funding support from the National Institutes of Health (GM084018 to A.R.). R. Z. greatly acknowledges the support of the Fundamental Research Funds for the Central University.

## References

[1] T. Schubeis, T. Le Marchand, L.B. Andreas, G. Pintacuda, 1H magic-angle spinning NMR evolves as a powerful new tool for membrane proteins, J. Magn. Reson., 287 (2018) 140–152.

[2] M. Tang, G. Comellas, C.M. Rienstra, Advanced Solid-State NMR Approaches for Structure Determination of Membrane Proteins and Amyloid Fibrils, Acc. Chem. Res, 46 (2013) 2080–2088.

[3] L.A. Baker, M. Baldus, Characterization of membrane protein function by solid-state NMR spectroscopy, Curr. Opin. Struct. Biol, 27 (2014) 48–55.

[4] S. Wang, V. Ladizhansky, Recent advances in magic angle spinning solid state NMR of membrane proteins, Prog. Nucl. Magn. Reson. Spectrosc, 82 (2014) 1–26.

[5] V.S. Mandala, J.K. Williams, M. Hong, Structure and Dynamics of Membrane Proteins from Solid-State NMR, Annu. Rev. Biophys. 47 (2018), 201–222.

[6] H.W. Spiess, 50th Anniversary perspective: the importance of NMR spectroscopy to macromolecular Science, Macromolecules, 50 (2017) 1761–1777.

[7] M.R. Hansen, R. Graf, H.W. Spiess, Interplay of structure and dynamics in functional macromolecular and supramolecular systems as revealed by magnetic resonance spectroscopy, Chem. Rev, 116 (2015) 1272–1308.

[8] K. Schmidt-Rohr, H.W. Spiess, Multidimensional solid-state NMR and polymers, Academic Press, London, 1994.

[9] K.H. Mroue, A. Viswan, N. Sinha, A. Ramamoorthy, Solid-State NMR Spectroscopy: The Magic Wand to View Bone at Nanoscopic Resolution, Annu. Rep. NMR. Spectrosc., 92(2017)365–413.

[10] M.J. Duer, The contribution of solid-state NMR spectroscopy to understanding biomineralization: Atomic and molecular structure of bone, J. Magn. Reson. 253 (2015) 98–110.

[11] Y.-Y. Hu, A. Rawal, K. Schmidt-Rohr, Strongly bound citrate stabilizes the apatite nanocrystals in bone, Proc. Natl. Acad. Sci. U. S. A. 107 (2010) 22425–22429.

[12] T. Theint, P.S. Nadaud, D. Aucoin, J.J. Helmus, S.P. Pondaven, K. Surewicz, W.K. Surewicz, C.P. Jaroniec, Species-dependent structural polymorphism of Y145Stop prion protein amyloid revealed by solid-state NMR spectroscopy, Nat. Comm. 8 (2017) 753.

[13] M. Lee, T. Wang, O.V. Makhlynets, Y. Wu, N.F. Polizzi, H. Wu, P.M. Gosavi, J. Stöhr, I.V. Korendovych, W.F. DeGrado, Zinc-binding structure of a catalytic amyloid from solid-state NMR, Proc. Natl. Acad. Sci. U. S. A. 114 (2017) 6191–6196.

[14] Z. Niu, W. Zhao, Z. Zhang, F. Xiao, X. Tang, J. Yang, The Molecular Structure of Alzheimer β-Amyloid Fibrils Formed in the Presence of Phospholipid Vesicles, Angew. Chem. Int. Ed. 53 (2014) 9294–9297.

[15] W. Qiang, W.-M. Yau, J.-X. Lu, J. Collinge, R. Tycko, Structural variation in amyloid-β fibrils from Alzheimer’s disease clinical subtypes, Nature, 541 (2017) 217.

[16] L. Lecoq, T. Wiegand, F.J. Rodriguez-Alvarez, R. Cadalbert, G.A. Herrera, L. del Pozo-Yauner, B.H. Meier, A. Böckmann, A Substantial Structural Conversion of the Native Monomer Leads to in-Register Parallel Amyloid Fibril Formation in Light-Chain Amyloidosis, Chembiochem, 20 (2019) 1027–1031.

[17] A. Mainz, J. Peschek, M. Stavropoulou, K.C. Back, B. Bardiaux, S. Asami, E. Prade, C. Peters, S. Weinkauf, J. Buchner, B. Reif, The chaperone αB-crystallin uses different interfaces to capture an amorphous and an amyloid client, Nat. Struct. Mol. Biol. 22 (2015) 898.

[18] Y. Gao, R. Zhang, W. Lv, Q. Liu, X. Wang, P. Sun, H.H. Winter, G. Xue, Critical Effect of Segmental Dynamics in Polybutadiene / Clay Nanocomposites Characterized by Solid State 1H NMR Spectroscopy, J. Phys. Chem. C. 118 (2014) 5606–5614.

[19] J. Brus, M. Urbanová, I. Kelnar, J. Kotek, A Solid-State NMR Study of Structure and Segmental Dynamics of Semicrystalline Elastomer-Toughened Nanocomposites, Macromolecules, 39 (2006) 5400–5409.

[20] C. Bonhomme, C. Gervais, D. Laurencin, Recent NMR developments applied to organic-inorganic materials, Prog. in Nucl. Magn. Reson. Spectrosc., 77 (2014) 1–48.

[21] Q. Dang, S. Lu, S. Yu, P. Sun, Z. Yuan, Silk Fibroin/Montmorillonite Nanocomposites: Effect of pH on the Conformational Transition and Clay Dispersion, Biomacromolecules, 11 (2010) 1796–1801.

[22] K. Takegoshi, S. Nakamura, T. Terao, 13C-1H dipolar-assisted rotational resonance in magic-angle spinning NMR, Chem. Phys. Lett., 344 (2001) 631–637.

[23] M. Shen, B. Hu, O. Lafon, J. Trébosc, Q. Chen, J.-P. Amoureux, Broadband finite-pulse radio-frequency-driven recoupling (fp-RFDR) with (XY8)41 super-cycling for homo-nuclear correlations in very high magnetic fields at fast and ultra-fast MAS frequencies, J. Magn. Reson., 223 (2012) 107–119.

[24] J.z.R. Lewandowski, G.l.D. Paëpe, M.T. Eddy, R.G. Griffin, 15N−15N Proton Assisted Recoupling in Magic Angle Spinning NMR, J. Am. Chem. Soc., 131 (2009) 5769–5776.

[25] Y. Ishii, 13C–13C dipolar recoupling under very fast magic angle spinning in solid-state nuclear magnetic resonance: Applications to distance measurements, spectral assignments, and high-throughput secondary-structure determination, J. Chem. Phys., 114 (2001) 8473–8483.

[26] Y. Jayasubba Reddy, V. Agarwal, A. Lesage, L. Emsley, K.V. Ramanathan, Heteronuclear proton double quantum-carbon single quantum scalar correlation in solids, J. Magn. Reson., 245 (2014) 31–37.

[27] D.H. Zhou, G. Shah, M. Cormos, C. Mullen, D. Sandoz, C.M. Rienstra, Proton-Detected Solid-State NMR Spectroscopy of Fully Protonated Proteins at 40 kHz Magic-Angle Spinning, J. Am. Chem. Soc., 129 (2007) 11791–11801.

[28] G.P. Holland, Q. Mou, J.L. Yarger, Determining hydrogen-bond interactions in spider silk with 1H-13C HETCOR fast MAS solid-state NMR and DFT proton chemical shift calculations, Chem. Commun., 49 (2013) 6680–6682.

[29] B.-J. Van Rossum, C. De Groot, V. Ladizhansky, S. Vega, H. De Groot, A method for measuring heteronuclear (1H− 13C) distances in high speed MAS NMR, J. Am. Chem. Soc., 122 (2000) 3465–3472.

[30] S.M. Althaus, K. Mao, J.A. Stringer, T. Kobayashi, M. Pruski, Indirectly detected heteronuclear correlation solid-state NMR spectroscopy of naturally abundant 15N nuclei, Solid State Nucl. Magn. Reson., 57–58 (2014) 17-21.

[31] K. Mao, M. Pruski, Directly and indirectly detected through-bond heteronuclear correlation solid-state NMR spectroscopy under fast MAS, J. Magn. Reson., 201 (2009) 165–174.

[32] M.D. Tuttle, G. Comellas, A.J. Nieuwkoop, D.J. Covell, D.A. Berthold, K.D. Kloepper, J.M. Courtney, J.K. Kim, A.M. Barclay, A. Kendall, Solid-state NMR structure of a pathogenic fibril of full-length human α-synuclein, Nat. Struct. Mol. Biol. 23 (2016) 409.

[33] V. Agarwal, S. Penzel, K. Szekely, R. Cadalbert, E. Testori, A. Oss, J. Past, A. Samoson, M. Ernst, A. Böckmann, De Novo 3D Structure Determination from Sub-milligram Protein Samples by Solid-State 100 kHz MAS NMR Spectroscopy, Angew. Chem. Int. Ed., 53 (2014) 12253–12256.

[34] A. Marchanka, B. Simon, G. Althoff-Ospelt, T. Carlomagno, RNA structure determination by solid-state NMR spectroscopy, Nature Commun., 6 (2015) 7024.

[35] J.-P. Demers, B. Habenstein, A. Loquet, S.K. Vasa, K. Giller, S. Becker, D. Baker, A. Lange, N.G. Sgourakis, High-resolution structure of the Shigella type-III secretion needle by solid-state NMR and cryo-electron microscopy, Nature Commun., 5 (2014) 4976.

[36] S. Wang, R.A. Munro, L. Shi, I. Kawamura, T. Okitsu, A. Wada, S.-Y. Kim, K.-H. Jung, L.S. Brown, V. Ladizhansky, Solid-state NMR spectroscopy structure determination of a lipid-embedded heptahelical membrane protein, Nat. Meth., 10 (2013) 1007–1012.

[37] S. Yan, C.L. Suiter, G. Hou, H. Zhang, T. Polenova, Probing structure and dynamics of protein assemblies by magic angle spinning NMR spectroscopy, Acc. Chem. Res., 46 (2013) 2047–2058.

[38] G. Hou, S. Yan, S. Sun, Y. Han, I.-J.L. Byeon, J. Ahn, J. Concel, A. Samoson, A.M. Gronenborn, T. Polenova, Spin Diffusion Driven by R-Symmetry Sequences: Applications to Homonuclear Correlation Spectroscopy in MAS NMR of Biological and Organic Solids, J. Am. Chem. Soc., 133 (2011) 3943–3953.

[39] V. Kurauskas, E. Crublet, P. Macek, R. Kerfah, D.F. Gauto, J. Boisbouvier, P. Schanda, Sensitive proton-detected solid-state NMR spectroscopy of large proteins with selective CH3 labelling: application to the 50S ribosome subunit, Chem. Commun., 52 (2016) 9558–9561.

[40] R. Zhang, K.H. Mroue, A. Ramamoorthy, Proton-Based Ultrafast Magic Angle Spinning Solid-State NMR Spectroscopy, Acc. Chem. Res., 50 (2017) 1105–1113.

[41] Y. Nishiyama, Fast magic-angle sample spinning solid-state NMR at 60-100 kHz for natural abundance samples, Solid State Nucl. Magn. Reson., 78 (2016) 24–36.

[42] R. Zhang, Y. Nishiyama, P. Sun, A. Ramamoorthy, Phase cycling schemes for finite-pulse-RFDR MAS solid state NMR experiments, J. Magn. Reson., 252 (2015) 55–66.

[43] A. Krajnc, T. Kos, N. Zabukovec Logar, G. Mali, A Simple NMR-Based Method for Studying the Spatial Distribution of Linkers within Mixed-Linker Metal–Organic Frameworks, Angew. Chem. Int. Ed., 54(2015) 10535–10538.

[44] A.J. Rossini, M.P. Hanrahan, M.M. Thuo, Rapid Acquisition of Wideline MAS Solid-state NMR Spectra with Fast MAS, Indirect Proton Detection, and Dipolar HMQC Pulse Sequences, Phys. Chem. Chem. Phys. 18(2016)25284–25295.

[45] P. Fricke, V. Chevelkov, M. Zinke, K. Giller, S. Becker, A. Lange, Backbone assignment of perdeuterated proteins by solid-state NMR using proton detection and ultrafast magic-angle spinning, Nat. Protoc., 12 (2017) 764.

[46] D. Mukhopadhyay, P.S. Nadaud, M.D. Shannon, C.P. Jaroniec, Rapid Quantitative Measurements of Paramagnetic Relaxation Enhancements in Cu(II)-Tagged Proteins by Proton-Detected Solid-State NMR Spectroscopy, J. Phys. Chem. Lett. 8 (2017)5871–5877.

[47] J. Tolchard, M.K. Pandey, M. Berbon, A. Noubhani, S.J. Saupe, Y. Nishiyama, B. Habenstein, A. Loquet, Detection of side-chain proton resonances of fully protonated biosolids in nano-litre volumes by magic angle spinning solid-state NMR, J. Biomol. NMR, 70(2018)177–185.

[48] L.B. Andreas, T. Le Marchand, K. Jaudzems, G. Pintacuda, High-resolution proton-detected NMR of proteins at very fast MAS, J. Magn. Reson., 253 (2015) 36–49.

[49] Y. Ishii, J.P. Yesinowski, R. Tycko, Sensitivity Enhancement in Solid-State 13C NMR of Synthetic Polymers and Biopolymers by 1H NMR Detection with High-Speed Magic Angle Spinning, J. Am. Chem. Soc., 123 (2001) 2921–2922.

[50] R. Bernd, Ultra-high resolution in MAS solid-state NMR of perdeuterated proteins: Implications for structure and dynamics, J. Magn. Reson., 216 (2012) 1–12.

[51] S.K. Vasa, P. Rovó, R. Linser, Protons as Versatile Reporters in Solid-State NMR Spectroscopy, Acc. Chem. Res., 51 (2018) 1386–1395.

[52] E.K. Paulson, C.R. Morcombe, V. Gaponenko, B. Dancheck, R.A. Byrd, K.W. Zilm, Sensitive High Resolution Inverse Detection NMR Spectroscopy of Proteins in the Solid State, J. Am. Chem. Soc., 125 (2003) 15831–15836.

[53] J. Schaefer, E. Stejskal, Carbon-13 nuclear magnetic resonance of polymers spinning at the magic angle, J. Am. Chem. Soc., 98 (1976) 1031–1032.

[54] A. Pines, M. Gibby, J. Waugh, Proton-enhanced NMR of dilute spins in solids, J. Chem. Phys., 59 (1973) 569–590.

[55] R. Zhang, K.H. Mroue, A. Ramamoorthy, Hybridizing cross-polarization with NOE or refocused-INEPT enhances the sensitivity of MAS NMR spectroscopy, J. Magn. Reson., 266 (2016) 59–66.

[56] A. Bielecki, A. Kolbert, M. Levitt, Frequency-switched pulse sequences: homonuclear decoupling and dilute spin NMR in solids, Chem. Phys. Lett., 155 (1989) 341–346.

[57] A. Bielecki, A. Kolbert, H. De Groot, R. Griffin, M. Levitt, Frequency-Switched Lee—Goldburg Sequences in Solids, Adv. Magn. Reson. 14(1990) 111–124.

[58] M. Lee, W.I. Goldburg, Nuclear-magnetic-resonance line narrowing by a rotating rf field, Phys. Rev., 140 (1965) A1261.

[59] M. Mehring, J. Waugh, Magic-angle NMR experiments in solids, Phys. Rev. B, 5 (1972) 3459.

[60] N.C. Nielsen, H. Bildso/e, H.J. Jakobsen, M.H. Levitt, Double-quantum homonuclear rotary resonance: Efficient dipolar recovery in magic-angle spinning nuclear magnetic resonance, J. Chem. Phys., 101 (1994) 1805–1812.

[61] W.G. Hu, K. Schmidt-Rohr, Polymer ultradrawability: the crucial role of α-relaxation chain mobility in the crystallites, Acta Polym., 50 (1999) 271–285.

[62] C. Tang, A. Inomata, Y. Sakai, H. Yokoyama, T. Miyoshi, K. Ito, Effects of Chemical Modification on the Molecular Dynamics of Complex Polyrotaxanes Investigated by Solid-State NMR, Macromolecules, 46 (2013) 6898–6907.

[63] X. Yuan, D. Sperger, E.J. Munson, Investigating Miscibility and Molecular Mobility of Nifedipine-PVP Amorphous Solid Dispersions Using Solid-State NMR Spectroscopy, Mol. Pharm., 11 (2014) 329–337.

[64] K.R. Mote, V. Agarwal, P.K. Madhu, Five decades of homonuclear dipolar decoupling in solid-state NMR: Status and outlook, Prog. Nucl. Magn. Reson. Spectrosc., 97 (2016) 1–39.

[65] P.K. Madhu, High-resolution solid-state NMR spectroscopy of protons with homonuclear dipolar decoupling schemes under magic-angle spinning, Solid State Nucl. Magn. Reson., 35 (2009) 2–11.

[66] A.M. Jordan, K. Kim, D. Soetrisno, J. Hannah, F.S. Bates, S.A. Jaffer, O. Lhost, C.W. Macosko, Role of Crystallization on Polyolefin Interfaces: An Improved Outlook for Polyolefin Blends, Macromolecules, 51(2018)2506–2516.

[67] K. Ishizuki, D. Aoki, R. Goseki, H. Otsuka, Multicolor Mechanochromic Polymer Blends That Can Discriminate between Stretching and Grinding, ACS Macro Letters, 7(2018) 556–560.

[68] S. Lu, R. Zhang, X. Wang, P. Sun, W. Lv, Q. Liu, N. Jia, Effect of PEO molecular weight on the miscibility and dynamics in epoxy/PEO blends, Euro. Phys. J. E, 38 (2015) 1–8.

[69] D. Tian, T. Li, R. Zhang, Q. Wu, T. Chen, P. Sun, A. Ramamoorthy, Conformations and Intermolecular Interactions in Cellulose/Silk Fibroin Blend Films: A Solid-State NMR Perspective, J. Phys. Chem. B, 121 (2017) 6108–6116.

[70] T.H. Chen, J.J. Zhu, B.H. Li, S.Y. Guo, Z.Y. Yuan, P.C. Sun, D.T. Ding, A.C. Shi, Exfoliation of organo-clay in telechelic liquid polybutadiene rubber, Macromolecules, 38 (2005) 4030–4033.

[71] T. Gopinath, S.E.D. Nelson, G. Veglia, 1H-detected MAS solid-state NMR experiments enable the simultaneous mapping of rigid and dynamic domains of membrane proteins, J. Magn. Reson., 285 (2017) 101–107.

[72] T. Gopinath, G. Veglia, 3D DUMAS: Simultaneous acquisition of three-dimensional magic angle spinning solid-state NMR experiments of proteins, J. Magn. Reson., 220 (2012) 79–84.

[73] T. Gopinath, G. Veglia, Dual Acquisition Magic-Angle Spinning Solid-State NMR-Spectroscopy: Simultaneous Acquisition of Multidimensional Spectra of Biomacromolecules, Angew. Chem. Int. Ed. 51(2012)2731–2735.

[74] K. Sharma, P.K. Madhu, K.R. Mote, A suite of pulse sequences based on multiple sequential acquisitions at one and two radiofrequency channels for solid-state magic-angle spinning NMR studies of proteins, J. Biomol. NMR, 65 (2016) 127–141.

[75] C. Martineau, F. Decker, F. Engelke, F. Taulelle, Parallelizing acquisitions of solid-state NMR spectra with multi-channel probe and multi-receivers: Applications to nanoporous solids, Solid State Nucl. Magn. Reson., 55–56 (2013) 48–53.

[76] Ē. Kupče, K.R. Mote, P.K. Madhu, Experiments with direct detection of multiple FIDs, J. Magn. Reson., 304 (2019) 16–34.

[77] R. Zhang, Y. Chen, N. Rodriguez-Hornedo, A. Ramamoorthy, Enhancing NMR Sensitivity of Natural-Abundance Low-γ Nuclei by Ultrafast Magic-Angle-Spinning Solid-State NMR Spectroscopy, ChemPhysChem, 17 (2016) 2962–2966.

[78] S.A. Kotler, J.R. Brender, S. Vivekanandan, Y. Suzuki, K. Yamamoto, M. Monette, J. Krishnamoorthy, P. Walsh, M. Cauble, M.M.B. Holl, E.N.G. Marsh, A. Ramamoorthy, High-resolution NMR characterization of low abundance oligomers of amyloid-β without purification, Sci. Rep., 5 (2015) 11811.

